# Imaging Intra- and Extracellular Conductivity using MR based Conductivity Tensor Imaging

**DOI:** 10.1101/2024.07.01.601471

**Authors:** Neha Rana, Nitish Katoch

**Affiliations:** Institute for Basic Sciences, Center for Soft and Living Matter, Ulsan 44919, Republic of Korea; Department of Chemical Engineering, Ulsan National Institute of Science and Technology, Ulsan 44919, Republic of Korea; Department of Biomedical Engineering, Kyung Hee University, Seoul 02447, Republic of Korea

**Keywords:** Electrical Conductivity, Image Reconstruction, Image Processing, Magnetic Resonance Imaging (MRI), Conductivity Tensor Imaging (CTI)

## Abstract

Imaging electrical conductivity may reveal relationships between biological tissues, cellular structures, and physiological processes. Biological tissues are primarily composed of major ions such as Na+ and K+, with varying concentrations and mobility within the cellular structures. These tissues consist of intracellular and extracellular fluids separated by cell membranes, and their electrical conductivity can be expressed as a function of ion concentration and mobility. This study introduces Conductivity Tensor Imaging (CTI) to independently reconstruct the electrical conductivity of intra- and extracellular compartments in biological tissues using MRI. We validated this method using a conductivity phantom with three compartments filled with electrolytes and/or giant vesicle suspensions. These vesicles mimic cell-like materials with thin insulating membranes, providing a realistic model for cellular structures. Measurements showed that high-frequency conductivity closely matched low-frequency conductivity in normal electrolytes. However, in the giant vesicle compartment, the conductivity of extracellular (EC) and intracellular (IC) regions correlated with cell volume fraction. In vivo human brain imaging using CTI revealed significant EC and IC conductivity variations across different brain regions, corresponding to underlying cellular compositions and structures. CTI introduces a novel MR contrast mechanism to distinctly measure IC and EC conductivities. Our findings highlight the potential of CTI to enhance our understanding of brain microstructure and its physiological processes through detailed conductivity mapping. This method signifies a notable advancement in non-invasive imaging, providing novel insights into the electrical properties of biological tissues and their implications for biophysical properties.

## Introduction

The electrical properties of biological tissues primarily reflect cellular structures, including cell density, volume fraction, and water content, as well as the concentration and mobility of major ions in both intra- and extracellular fluids. Factors such as cell size, shape, fluid composition, temperature, hydration level, and viscosity significantly influence the mobility of charged carriers [1]. The structural features of cells also influence mobility. For instance, when elongated cells are aligned in a specific direction, the movement of ions in the extracellular fluid is hindered. This phenomenon demonstrates the property of tissue anisotropy [2-4]. The electrical conductivity of a biological tissue is determined not only by the concentrations and mobility of major ions in the extracellular and intracellular fluids, but also by its structural properties such as the extracellular volume fraction, extracellular matrix materials and cellular structure [5, 6]. Brain tissues, such as white matter and gray matter, are composed of neurons and glial cells surrounded by extracellular fluid and matrix materials [7]. These cells exhibit distinct ionic configurations and structural properties crucial for electrical conductivity, owing to their specialized lipid bilayer membranes enveloping various organelles in an intracellular fluid [4].

Cross-sectional imaging of electrical conductivity distributions in the human body has been pursued for over 30 years [8]. Techniques such as MR-based electrical impedance tomography (MREIT) and magnetic resonance electrical properties tomography (MREPT) have been developed. MREIT reconstructs low-frequency isotropic conductivity images by injecting low-frequency currents and measuring induced magnetic flux density distributions using an MRI scanner [9]. MREPT produces conductivity and permittivity images at 128 MHz using a 3 T MRI scanner [10]. Based on a B1 mapping technique, it measures the effects of eddy currents induced by RF pulses. At the Larmor frequency of 128 MHz at 3T, however, useful low-frequency conductivity contrast information disappears [10, 11]. For most physiological events and pathological states observable below 1 kHz, a new noninvasive in vivo imaging method is needed to quantitatively visualize low-frequency conductivity tensor distributions inside the human body [12]. In oncology, MREPT has been widely used, correlating with in-vitro and histological findings [13]. Tumor conductivity measurement has proven effective in distinguishing patients with grade II, III, and IV gliomas during 3T clinical scans [14]. Recently, an electrodeless conductivity tensor imaging (CTI) technique was proposed to produce low-frequency conductivity tensor images using an MRI scanner [15, 16]. Previous studies have established the recovery of extracellular conductivity from high-frequency measurements using MREPT [15, 16], but distinguishing conductivity for both intra- and extracellular compartments has not been achieved. In this study, we introduce a novel CTI method designed to differentiate high frequency conductivity (HFC) into extracellular (EC) and intracellular conductivity (IC). Our approach leverages MR-based conductivity tensor imaging (CTI), uniquely suited for probing the ionic microenvironment within intra- and extracellular spaces. We validate our method using both a conductivity phantom and human brain, demonstrating its feasibility in characterizing critical microenvironmental properties of conductivity.

This paper presents a new approach for reconstructing the conductivity of intra- and extracellular spaces in the human brain using two MR imaging scans: B1 phase maps for HFC reconstruction and diffusion tensor imaging (DTI) for distinguishing spaces and their respective diffusion coefficients. Initial validation employed a cell-like phantom comprising unilamellar giant vesicles (GVS), engineered to mimic cellular properties with thin insulating membranes that replicate both extracellular and intracellular architectures. GVS, due to their size and structure similar to biological cells, feature closed bilayer membranes. They emulate eukaryotic cells with ‘vesicle-in-vesicle’ configurations, forming droplet-like structures through gentle hydration, electroformation, or microfluidic techniques. Ionic concentrations were meticulously controlled within intra- and extracellular domains. Furthermore, the proposed method was applied to the human brain to measure in vivo tissue conductivities. This study reports the results of validating in vitro intra- and extracellular conductivity, comparing measurements from giant vesicle phantoms with those obtained using impedance analyzers. Notably, this study represents the first investigation into intra- and extracellular conductivity in the human brain.

## Materials and Methods

### Phantom preparations

Giant vesicles (GVS) resemble cells with thin insulating membrane and are prepared using the reversal phase method following process described in Mosche et al [17, 18]. Giant vesicles dispersed in an electrolyte were prepared following a detailed protocol available at [5]. Initially, 2 mL of phospholipids (Avanti Polar Lipids, Alabaster, AL, https://avantilipids.com) were dissolved in chloroform at a concentration of 30 mg/mL under an argon atmosphere in a round-bottom flask. This flask was then connected to a rotary evaporator (N-1300V-W, EYELA, Tokyo, Japan) to remove the organic solvent. The evaporation process involved two phases: first, the solution was subjected to 47°C under vacuum with a nitrogen trap for 20 minutes at 10 rpm. Then, the rotation speed was increased to 60 rpm for an additional 20 minutes. During this evaporation period, the phospholipids self-assembled into giant vesicles. After evaporation, the aqueous solution containing the giant vesicles was centrifuged at 1500 rpm for 10 minutes. The prepared suspension and microscopic images of the vesicles are shown in Figure 1A. The resulting giant vesicle suspensions were visually inspected under a microscope, with volume fractions estimated to range between 80% and 90%. This protocol ensures the effective preparation and isolation of giant vesicles for further experimental use. The phantom used in the experiments consists of three compartments, as illustrated in Figure 2A. The phantom has a diameter of 40 mm and a height of 75 mm. It contains two electrolytes with different ion concentrations: Electrolyte #1 is a NaCl solution at 7.5 g/L, and Electrolyte #2 is a mixture of 3.5 g/L NaCl and 1 g/L CuSO_4_. The giant vesicle suspension (GVS), with an average size of 13 ± 4.7 µm, was filled with Electrolyte #1 and suspended within the same electrolyte. The density of the giant vesicles was verified through visual observation, ensuring an accurate assessment of their distribution within the phantom. To validate the CTI results, the conductivity spectra of GVS were directly measured using an impedance analyzer (SI1260A, METEK, United Kingdom) with a four-electrode setup, as shown in Figure 1B.

**Figure 1.**
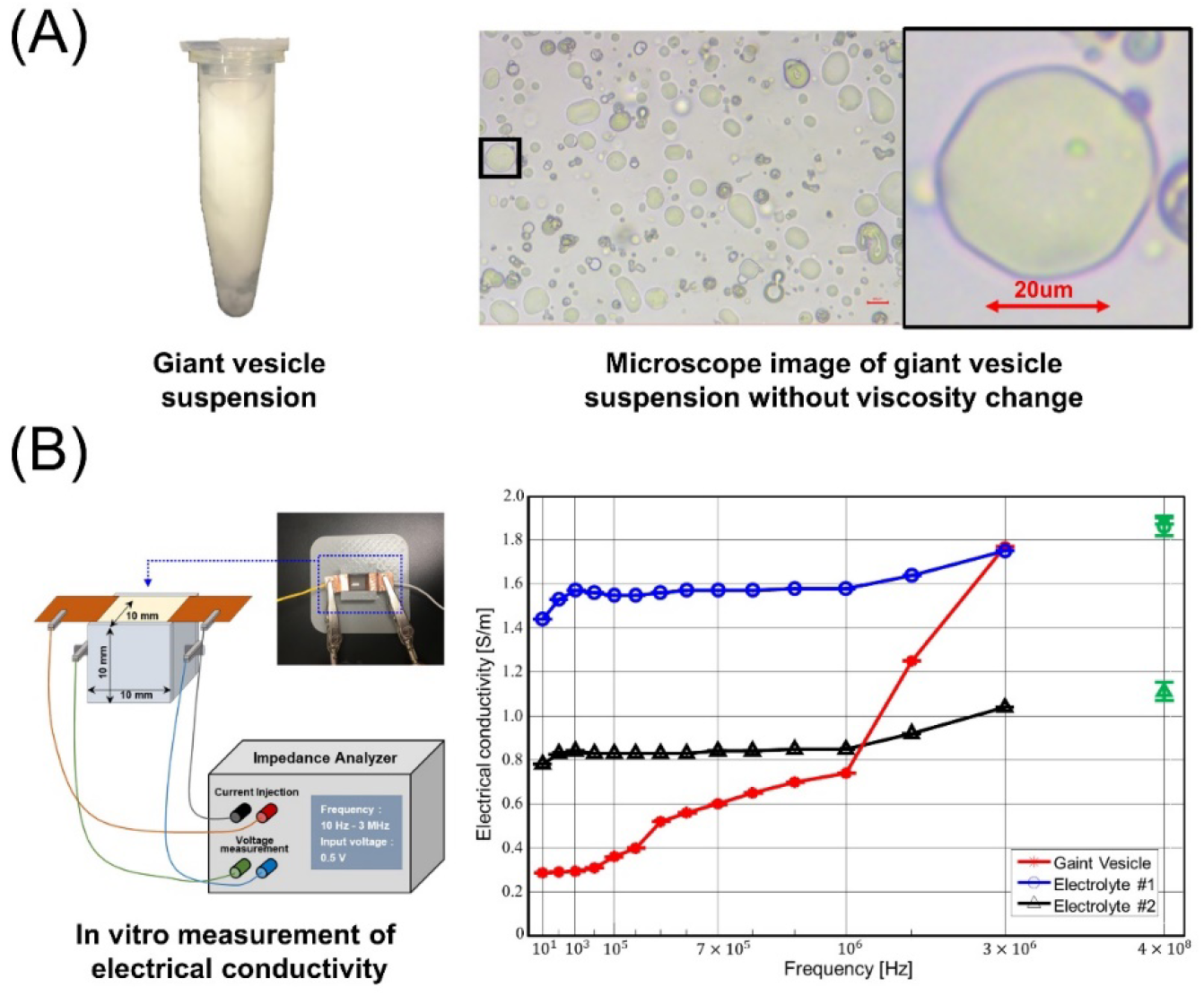
(A) Microscopic images of giant vesicles and the suspension medium in which these vesicles were placed. (B) In-vitro setup for measuring the conductivity of giant vesicles, along with the conductivity spectra of the phantom used for validation studies (Partially reproduced from Katoch et al [16]).

**Figure 2.**
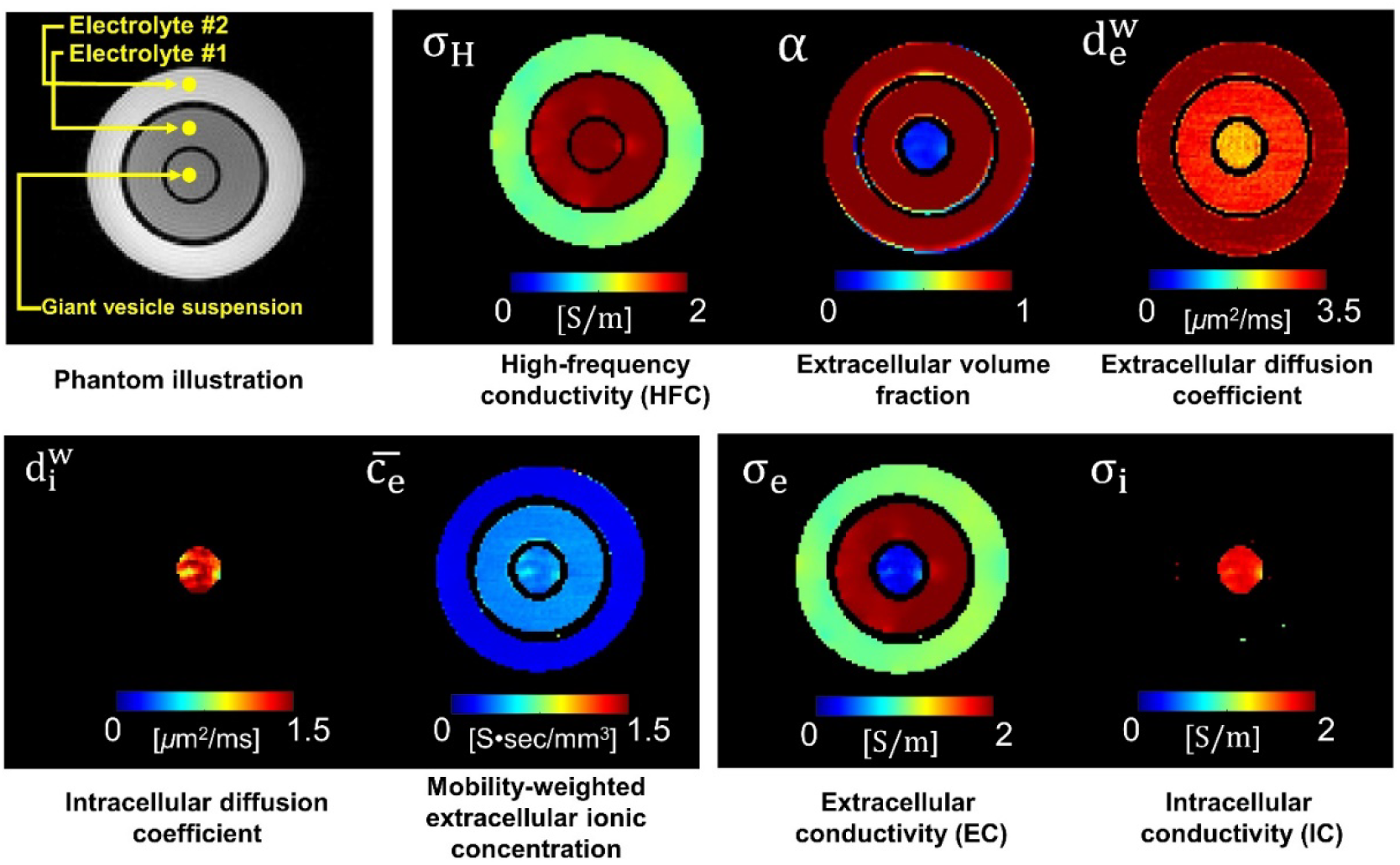
Images of CTI parameters for conductivity phantoms with giant vesicle suspensions. The parameters displayed include high-frequency conductivity (σ_H_), extracellular volume fraction (α), extracellular water diffusion Coefficient 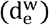, intracellular water diffusion coefficient 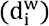 and mobility-weighted extracellular ionic concentrations 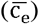. Additionally, extracellular conductivity (σ_e_) and intracellular conductivity (σ_i_), referred as EC and IC, respectively.

### Reconstruction algorithm: Intra- and Extracellular conductivity

The electrical conductivity of biological tissues can be expressed in terms of the ionic concentration and mobility of ions within intra- and extracellular spaces. These ions are the primary charge carriers in biological fluids [1, 4]. By applying the Einstein relation, which related the diffusivity and mobility of these charge carriers, the ionic mobility m_j_ of the jth charge carrier in both intra and extracellular spaces can be expressed as follows:

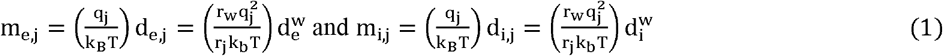

Where q_j_ = 1.6×10^−19^C charge of electron, k_B_ is the Boltzman constant, r_w_ and r_j_ are the Stoke’s radius of jth particle, T is the Absolute temperature, 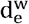 and 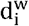 are the diffusion coefficients of water molecules in extracellular and intracellular compartments, respectively. Assuming that the reference charge carrier and water molecule exist in the same microscopic environment, we set 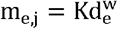 and 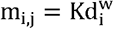.

Using the relation between the electrical conductivity and diffusion coefficient, the total tissue conductivity (σ_TTC_) in the biological tissues can be represented as:

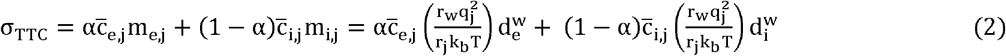

Where α is volume fraction of extracellular space and 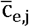 and 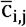 are the ionic concentrations in the extracellular and intracellular spaces, respectively. When utilizing MR-EPT, conductivity can be expressed as σ_TT_, as high frequency current (>1Khz) passes through thin, insulating cell membranes. Consequently, the measured high frequency conductivity σ_H_ = σ_TT_ has embedded information of conductivity.

Furthermore, assuming for some constant K ion concentration ratio 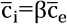, we can express the ion concentration in the extracellular space as:

Using equation 2, σ_H_ = σ_TTC_ and 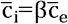, the ion concentration at extracellular space 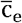 can be expressed as:

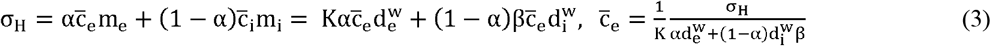

Low-frequency conductivity (σ_L_) is given by:

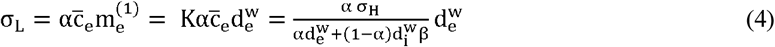

Where σ_L_ is low-frequency conductivity, derived from 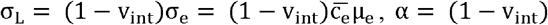 is extracellular volume fraction, β is ion concentration ratio and 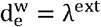 and 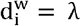 are the extra- and intracellular diffusion coefficients, respectively. σ_H_ is high-frequency conductivity (HFC), also referred to as total tissue conductivity.

To disentangle intra- and extracellular conductivity from σ_H_ and σ_L_ following equation can be used:

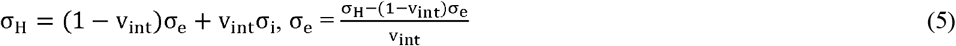

Assuming σ_e_ = σ_L_, intracellular conductivity (σ_i_) can be calculated. This approach enables the determination of both intra- and extracellular conductivities by using the measurements of high-frequency and low-frequency conductivities using CTI method. The reconstruction of 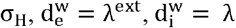 leveraging the methods described by Gurler et al. [19], Katoch et al. [16] and Jahng et al. [20].

*Simplified reconstruction steps:*

1. Measure high-frequency conductivity (σ_H_): σ_H_ = σ_TTC_
2. Calculate mobility-weighted effective extracellular ion concentration 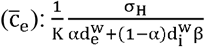
3. Determine extracellular conductivity(σ_e_): σ_e_ = σ_L_
4. Calculate intracellular conductivity 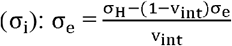

### MRI acquisition: CTI Experiment of Conductivity Phantom with Giant Vesicle Suspension

A 9.4 T research MRI scanner (Agilent Technologies, USA) equipped with a single-channel mouse body coil was used for this study. High-frequency conductivity images were acquired using a multi-echo spin-echo sequence with an isotropic voxel resolution of 0.5 mm. The imaging parameters were set as follows: TR/TE = 2200/22 ms, number of signal acquisitions = 5, field-of-view (FOV) = 65×65 mm^2^, slice thickness = 0.5 mm, flip angle = 90°, and image matrix = 128×128. Six echoes were acquired with a total scan time of 23 min.

Separately, diffusion-weighted MR imaging was separately performed using the single-shot spin-echo echo-planar-imaging sequence. The imaging parameters were TR/TE = 2000/70 ms, number of signal acquisitions = 2, FOV = 65×65 mm^2^, slice thickness = 0.5 mm, flip angle = 90°, and image matrix size = 128×128. The number of directions of the diffusion-weighting gradients was 30 with b-values of 50, 150, 300, 500, 700, 1000, 1400, 1800, 2200, 2600, 3000, 3600, 4000, 4500, and 5000 s/mm^2^. The phantom data used in this study is available at iirc.khu.ac.kr/software. Detailed protocols can be referenced from the work of Katoch et al. [5, 16].

### MRI acquisition: In-vivo human brain

The imaging experiments were performed using a 3T MRI scanner (Ingenia,, Philips Healthcare, Netherlands) equipped with a 32-channel head coil. The CTI method necessitates two separate MR imaging scans, with the parameters as follows:

#### High-frequency conductivity (HFC)

a balanced fast field echo (bFFE) sequence with TR/TE = 3/1.53 ms, resolution 2× 2×2 mm^3^, FOV 240×240 mm^2^ (sagittal slices). Non-selective RF pulses with FA = 25° and scan duration of 2 min 24 secs.

#### Microstructure imaging

This involved a multi-shell diffusion tensor imaging (DTI) acquired using a single-shot spin-echo echo-planar imaging (SS-SE-EPI) pulse sequence. The imaging parameters included b-shells with values of 500 (15 directions) 1000 (15 directions) and 2000 (30 directions) s/mm^2^ were performed. TR/TE=2700/63 ms, voxel size 2×2×2.5 mm^3^. The scan was accelerated using a SENSE factor of 3, resulting in a total scan time of 3 minutes. An additional T1 structural scan with resolution of 1 × 1 × 1 mm^3^ was performed to facilitate the segmentation of brain tissues. The images of intra- and extracellular conductivities were reconstructed using the CTI formula as described in Equation 1-5. The images were co-registered to magnitude of bFFE datasets.

## Results and Discussion

### Conductivity Measurements in Extracellular and Intracellular Spaces of Giant Vesicles

Fig. 2 shows the CTI parameter of high-frequency conductivity (σ_H_), extracellular volume fraction (α), extracellular water diffusion coefficient 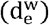, intracellular water diffusion coefficient 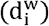 and mobility-weighted effective extracellular ion concentration 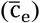. In phantom, σ_H_ images displayed higher contrast for Electrolyte #1 and the giant vesicle suspension compared to Electrolyte #2. This discrepancy is attributed to the differing ion concentrations between the two electrolytes. However, when the same Electrolyte #1 was used inside the giant vesicles, no contrast difference was observed in σ_H_. The measured parameter values in the phantom are presented in Table 1.

**Table 1.**
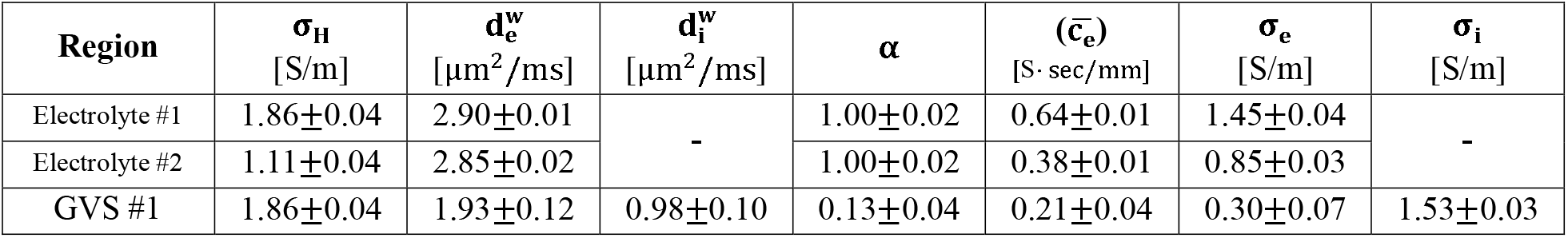
Mean and standard deviation values of intracellular and extracellular compartments and the intermediate variables used for reconstruction using the CTI method. The mean ± SD values were calculated from all pixels within each region.

Given that the same Electrolyte #1 was used both inside and outside the giant vesicles, the measured values of 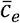 were found to be similar in both compartments, as shown in Figure 2 and Table 1. We also compared the conductivity values calculated using the equation 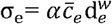 with those measured by an impedance analyzer in the giant vesicle phantom, observing a relative error ranging from 1.1% to 9.2%. In this study, the intracellular space in the electrolytic region of phantom was nonexistent, so the parameter *β* was considered as 0. When the same electrolyte was used both inside and outside the giant vesicles, the *β* was taken as 1. The reconstructed values of σ_i_ in Electrolytes #1 and #2 were close to zero, as there were no cells present, resulting in measured σ_H_ equating to σ_e_.

In the giant vesicle region, where cell membranes are present, high-frequency conductivity was not influenced by the cell membranes. However, the conductivity of the extracellular space was reduced due to the volume fraction (Table 1). These findings highlight the role of cell density in reducing extracellular conductivity, while conductivity at higher frequencies remains unaffected by the cell membrane. where cell membranes are present, high-frequency conductivity was not influenced by the cell membranes.

Using impedance analyzer (see figure 1), we also measured the *in-vitro* values of conductivity from samples used in phantom preparation. Focusing only on giant vesicles, the high-frequency conductivity (HFC) of the giant vesicle suspension was measured at 1.89 S/m at 400 MHz using the CTI experiment. This value is comparable to the 1.77 S/m measured at 3 MHz using an impedance analyzer (Figure 1B). The CTI method measured the extracellular conductivity (EC) at 0.29 S/m, which is within the error of 1.7%. The value of intracellular conductivity (IC) also correlates well with the amount of intracellular spaces (87%). Assuming the conductivity at 10 Hz approximates the DC conductivity, the measured HFC is weighted sum of IC and EC.

### Intra- and extracellular conductivity of human brain

Figure 3A shows the reconstructed high-frequency conductivity (HFC) separated into extracellular (σ_e_) and intracellular (σ_i_) conductivity, as illustrated in Figures 3F and 3G. Figures 3B-E display the microstructure parameters. The mean ± standard deviation values of these CTI parameters are detailed in Table 2. The conductivity values in Table 2 exhibit both subject and position dependency. For cerebrospinal fluid (CSF), the EC and HFC values were similar and close to zero, reflecting the lack of cells in CSF. In white matter (WM), the EC values were low, consistent with WM’s composition of tightly packed fiber bundles (axons) and reduced extracellular space. Consequently, the IC in WM is higher due to the denser intracellular environment. The conductivity values measured in HFC = EC were comparable to those found in the literature [16, 21, 22].

**Table 2.** The measured high frequency conductivity (*σ*_*H*_), extracellular conductivity *σ*_*e*_) and intracellular conductivity (*σ*_*i*_). The mean ± SD values were computed from regions in the brain.

| Region | $\sigma_H$ | $\sigma_e$ | $\sigma_i$ |
| --- | --- | --- | --- |

**Table 2.** The measured high frequency conductivity ( $\sigma_H$ ), extracellular conductivity ( $\sigma_e$ ) and intracellular conductivity ( $\sigma_i$ ). The mean ± SD values were computed from regions in the brain.
|  | [S/m] | [S/m] | [S/m] |
| --- | --- | --- | --- |
| CSF | $2.03 \pm 0.12$ | $2.01 \pm 0.11$ | $0.08 \pm 0.01$ |
| WM | $0.60 \pm 0.08$ | $0.20 \pm 0.04$ | $0.41 \pm 0.07$ |
| GM | $0.72 \pm 0.11$ | $0.52 \pm 0.09$ | $0.21 \pm 0.09$ |

**Figure 3.**
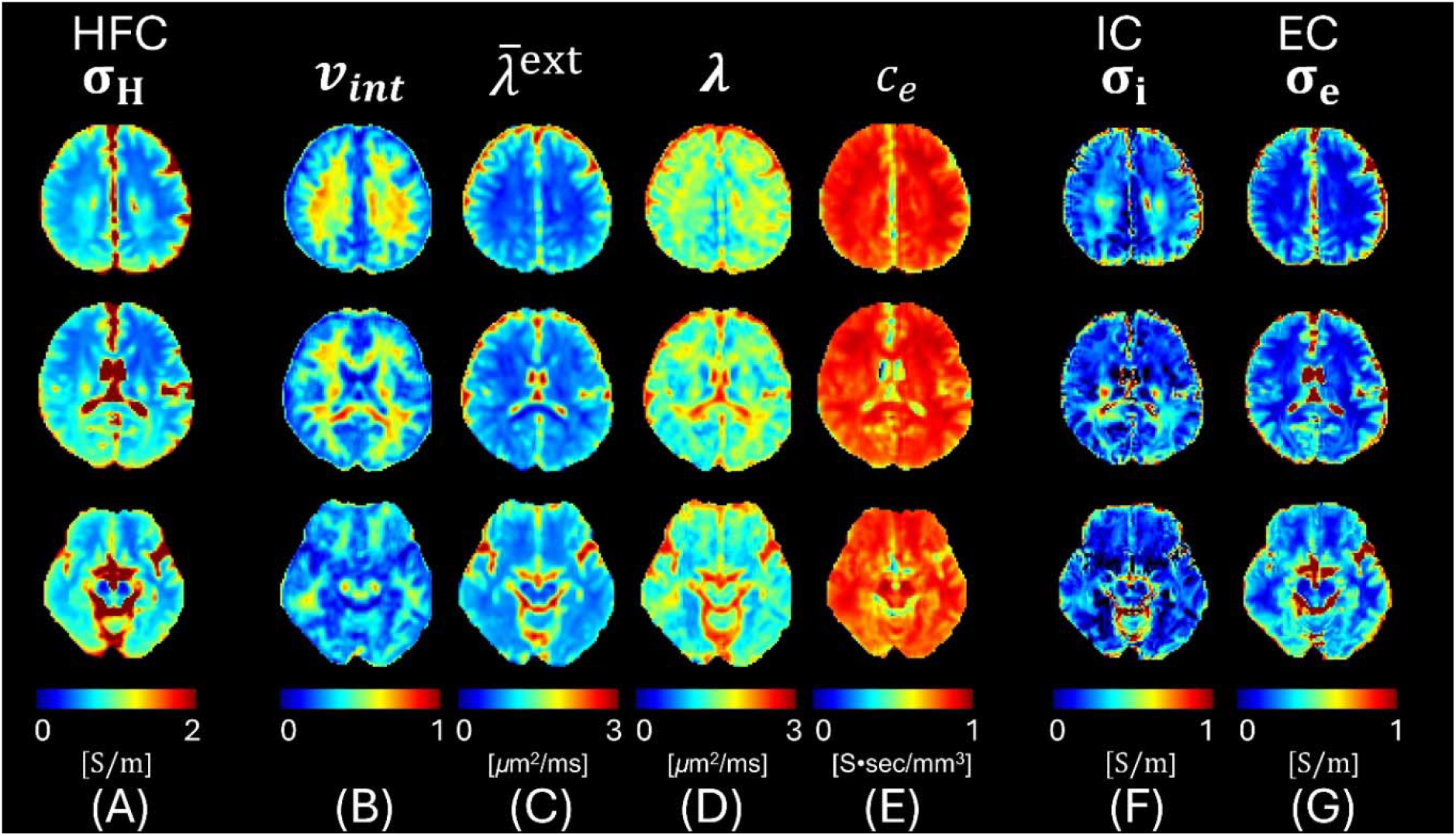
Conductivity tensor images of the human brain. (A) Reconstructed images of high-frequency conductivity (σ_H_), (B-E) Intermediate images of CTI. (F-G) Images of intra- and extracellular conductivity from three different slice positions.

WM predominantly consists of axons and oligodendrocytes. The flow of ions and their concentration in WM are influenced by the myelinated axons, where the myelin sheath restricts ion flow, leading to lower EC but higher IC due to the concentrated intracellular environment. In Table II, the gray matter conductivity values were between 0.20 and 0.30 S/m, which are close to the values of 0.24 to 0.29 S/m measured from in vivo DT-MREIT experiments of human subjects by using injection currents and also previously provided values [16, 22]. In gray matter (GM), similar trends in EC and IC were observed. GM contains a higher density of neuronal cell bodies and a lower density of glial cells compared to other regions, resulting in a lower IC. The flow of ions in GM is more dynamic due to the higher concentration of synapses and neural activity, leading to more balanced EC and IC values. Overall, the differences in EC and IC values across various brain regions align with the known cellular composition and density differences between CSF, WM, and GM. The ion flow and concentration vary significantly among these regions due to their distinct cellular structures: CSF lacks cells entirely, WM is rich in myelinated axons with restricted extracellular ion flow, and GM has dynamic ion flow due to its high neural activity and synaptic density.

The results from the human experiment supported the findings from the phantom experiment. The observed trends in conductivity in different brain regions correlate with the controlled conditions of the phantom, demonstrating the reliability and applicability of the CTI method in both *in-vitro* and *in-vivo* studies.

## Conclusion

In this study, we employed an electrodeless CTI method to visualize the distribution of intra- and extracellular conductivity. Through a phantom experiment, where we precisely controlled the concentrations of the ions, we validated that the proposed method can accurately extract ion-concentration information. Additionally, by incorporating cell-like material, we confirmed that the measured ion concentration originated from the extracellular space. The measured intracellular conductivity corroborated well with the cell density. The reconstructed parameters clearly demonstrated the effects of ionic concentration and mobility on electrical conductivity. Our *in vivo* measurements of human brain conductivity indicated significant variations in EC and IC across different regions, corresponding to the underlying cellular composition and structure. Specifically, the differences in conductivity values between cerebrospinal fluid, white matter, and gray matter align with the known cellular structures and densities in these regions. The CTI method can be easily implemented in clinical MRI scanners without the need for additional hardware, highlighting its feasibility and potential for broader clinical applications. Further research is essential to validate the potential clinical applications of these CTI methods. This includes exploring their use in characterizing tumors and Alzheimer’s disease, EEG source imaging, and planning treatments involving electrical stimulation.

## ACKNOWLEGEMENT

We would like to acknowledge the support from Impedance Imaging Research Center, Kyung Hee University. This publication fulfills the requirements for graduation.

## DATA AVAILABILITY

The presented data and codes used in the study is available at http://iirc.khu.ac.kr/toolbox.html

